# Sleep deprivation exacerbates seizures and diminishes GABAergic tonic inhibition

**DOI:** 10.1101/2021.03.06.434210

**Authors:** Sai Surthi Konduru, Yuzhen Pan, Eli Wallace, Jesse A Pfammatter, Mathew V. Jones, Rama K. Maganti

## Abstract

Patients with epilepsy report that sleep deprivation is a common trigger for breakthrough seizures. The basic mechanism of this phenomenon is unknown. In the Kv1.1^-/-^ mouse model of epilepsy, daily sleep deprivation indeed exacerbated seizures though these effects were lost after the 3^rd^ day. Sleep deprivation also accelerated mortality in ~52% of Kv1.1^-/-^ mice, not observed in controls. Voltage-clamp experiments on the day after recovery from sleep deprivation showed reductions in GABAergic tonic inhibition in dentate granule cells both in Kv1.1^-/-^ and wild-type mice. Our results suggest that sleep deprivation is detrimental to seizures and survival, possibly due to reductions in GABAergic tonic inhibition.

## Introduction

Sleep and epilepsy have a bidirectional relationship where seizures have sleep-wake patterns in various epilepsies and in turn, seizures disrupt sleep. Sleep deprivation (SD) is one of the most common triggers for breakthrough seizures^1^. SD is often used to activate diagnostic EEGs^2^ and in epilepsy monitoring units to trigger seizures. Some patient self-report studies showed that SD increases risk of breakthrough seizures, but a randomized controlled trial in an epilepsy-monitoring unit did not^3-4^. Regardless, the basic mechanisms underlying SD-induced seizure exacerbation are unknown.

In the Kv1.1 mouse model, homozygous mice with a functional mutation in the *Kcna1* gene that encodes the α-subunit of voltage-gated potassium channel^5^ have: a) spontaneous seizures with a sleep-wake or circadian pattern^6^, b) disrupted sleep and circadian rhythms^7^, c) progressive decline in time spent “resting” as mortality appoaches^8^ and d) premature mortality due to status epilepticus or SUDEP that occurs between ~P45-70^8^. In this model of temporal lobe epilepsy, we hypothesized that SD exacerbates seizures due to changes in GABAergic inhibition in the dentate gyrus.

## Methods

All procedures were performed after approval by the Institutional Animal Care and Use Committee (IACUC). Heterozygous mice (C3HeB.129S7-Kcna1tm1Tem/J, Jackson Labs) were interbred and appropriate genotyping procedures were performed to identify homozygous mice.

### Animal surgery

EEG electrode implantation was performed at P34-36 in all mice as previously described^6^. Briefly, mice were anesthetized with isoflurane (5% induction, 1-2% maintenance) and three stainless steel screw electrodes were implanted for EEG (bregma +1.5 mm and 1 mm right, bregma −3 mm and 1 mm left, and lambda −1 mm at midline) and two stainless steel braided wires in the nuchal muscles for EMG recording. After recovery, mice were transferred into individual recording chambers and allowed a ~24-hour acclimation period. Video EEG with EMG was acquired with an XLTek amplifier (XLTEK, USA) sampled at 1024 Hz.

### Sleep deprivation procedure

SD was performed for 4-hours a day x 5 days starting at lights-on (6:30 am) using gentle handling techniques (presenting novel objects for exploration or by gentle stroking with a soft paintbrush), starting at ~P37. EEG was recorded during a baseline day, during the 5-days of SD (n=19) and during one recovery day. Control animals (n=7) had seven continuous days of EEG recording with no SD. Two independent reviewers, blinded to treatment, manually analyzed EEG data for seizures (Figure 1A).

**Figure 1:**
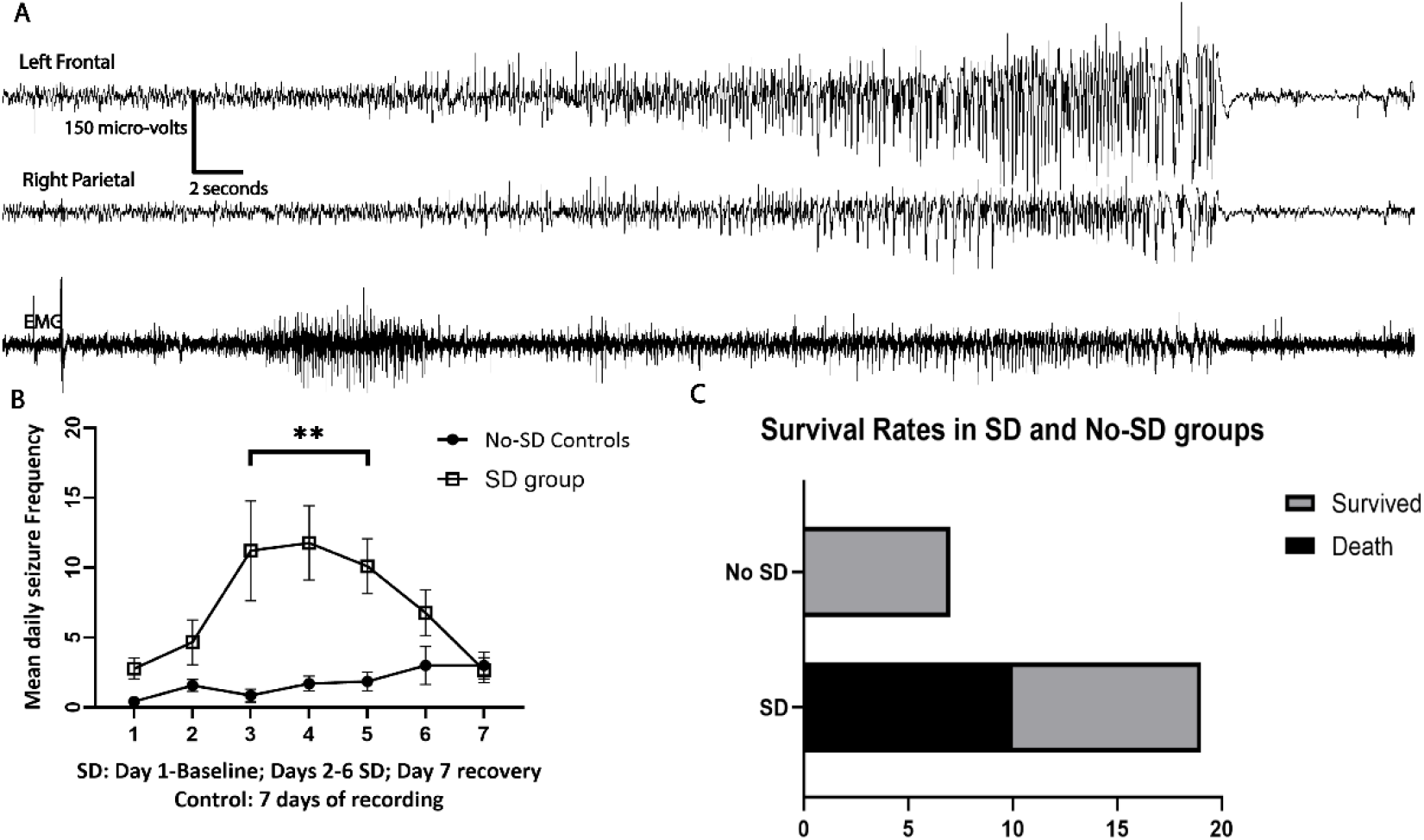
**Panel A**: An example of a seizure is shown from a Kv1.1^-/-^ mouse. Each mouse had a Right frontal and Left parietal EEG and an EMG channel shown. Seizures consisted of buildup of rhythmic discharges in frontal and parietal EEG lasting 20-60 seconds. **Panel B**: Mean daily seizure frequency (±SEM) between SD and non-SD controls across the 7 days was 2.5±0.5 vs 0.33±0.21 for day1; 3.5±1.11 vs 1.66±0.49 for day 2; 8.21±2.51 vs 1±0.51 on day 3; 10.92±2.36 vs 1.5±0.56 on day 4; 10.42±2.27 vs 1.33±0.49 on day 5; 6.777±1.36 vs 2.83±1.60 on day 6 and 2.66±0.89 vs 3.16±1.1 on day 7 with a statistically significant interaction between day of recording and SD treatment (Two-way ANOVA: F(6, 84)=3.03; *p*<0.001). Post-hoc Tukey tests showed SD group had significantly higher seizure frequency on Days 2, 3 and 4 of SD compared to baseline or recovery day (*p*<0.05**). No difference was seen in non-SD group across days of recording. **Panel C**: There was ~52% (10/19) mortality in the SD group where they died of status epilepticus during window of SD whereas 100% of non-SD group (7/7) survived the 7-day recording (*p*=0.02 Fisher’s exact test).

### Electrophysiology recordings

In a separate group of Kv1.1^-/-^ with and without SD (n=6 each) and wild-type mice that had SD (n=5), hippocampal slices were obtained and patch clamp electrophysiology was performed on the recovery day (SD group) or day-7 (non-SD) according to methods described previously^9^. Briefly, hippocampal slices were submerged in aCSF in a recording chamber. DG granular cells were visually identified with an IR-DIC microscope. Recordings in the voltage-clamp configuration (−70 mV) with a Axopatch 200B amplifier (Axon Instruments; Foster City, CA), low pass filtered at 2 kHz, and digitized at 10 kHz with a Digidata 1440A (Molecular Devices, San Jose, CA) were acquired with pCLAMP 10 (Molecular Devices, San Jose, CA). Borosilicate patch pipettes filled with KCl intracellular solution with a resistance of 2-5 MΩ when submerged in aCSF, were used to isolate GABAergic currents in a base solution containing 100 nM tetrodotoxin (TTX), 25μM (2R)-amino-5-phosphonovaleric acid (AP-5) and 10 μM 6-cyano-7-nitroquinoxaline-2,3-dione (CNQX). To measure GTI, 6-minute recordings were acquired in the base solution, followed by perfusion with 100 μM Bicuculline Methiodide(BIC), a GABA_A_ receptor antagonist used at a concentration that blocks all GABA_A_ receptors. Miniature IPSCs (mIPSCs) were detected using the MiniAnalysis program (Synaptosoft, Decatur, GA).

### Analysis

Interactions between SD-treatment and seizures across time was compared using two-way ANOVA with treatment group as a fixed variable and seizure frequency as a random variable, with post-hoc Tukey test for multiple comparisons. Differences in survival rates in SD and non-SD groups were compared using a Fisher’s exact test. Mean tonic currents (±SD) were calculated from all point amplitude histograms of 60-second segments of steady-state voltage-clamp traces, to which a Gaussian function was fit to the outward part of the current to avoid contamination by mIPSC currents. The magnitude of tonic current was calculated as the difference in fitted means between the baseline current and the current in the presence of BIC. Current density (pA/pF) was calculated by dividing the tonic current by cell capacitance. Differences in tonic current, current density, mIPSC amplitude and frequency were compared using one-way ANOVA with post hoc Tukey test.

## Results

### Sleep deprivation exacerbated seizure frequency

During the EEG recording in the SD group, 9/19 Kv1.1^-/-^ mice survived all 7 days and those that survived until day 4 of SD (n=14) were included in seizure analysis. Daily seizure frequency varied from 0-10 among non-SD control animals whereas it was 0-28 in the SD group. Seizure frequency increased in the SD group starting on the 2^nd^ day of SD, before declining after the 4^th^ day of SD (Figure 1B). No difference in seizure frequency across the 7-days of recording was seen for the non-SD group (*p*>0.05). No within group differences were seen for each day of the recording (*p*=0.23).

### Sleep deprivation enhanced mortality due to status epilepticus

During SD between ~P37 and 44, 10/19 (~52%) went into status epilepticus and all 10 died. Status epilepticus occurred anywhere from day 1 of SD to day 4 of SD, lasting 1-24 hours before an animal died. No mortality was seen among the non-SD or wild type controls (*p*=0.02, Figure 1C).

### Sleep deprivation diminished the magnitude of GABAergic tonic currents in dentate granule cells

During whole-cell voltage clamp (−70 mV) recordings, addition of the GABA_A_ receptor antagonist bicuculline to the perfusion solution resulted in blockade of mIPSCs as well as a reduction in holding current, signifying that GABA_A_ receptors contribute tonically to the resting conductance (Figure 2A). Both the tonic current magnitude and density diminished in sleep-deprived Kv1.1^-/-^ and wild types. Post-hoc Tukey tests showed that the Kv1.1^-/-^ with no-SD group had higher mean tonic current and current density than epileptic or wild-type animals that had SD (*p*<0.05) (Figure 2 B&C). No differences were also seen in mean mIPSC frequency or amplitude between SD and non-SD groups (*p*>0.05) (Figure 2 D&E). Findings suggest that SD impaired GABAergic tonic but not phasic inhibition.

**Figure 2:**
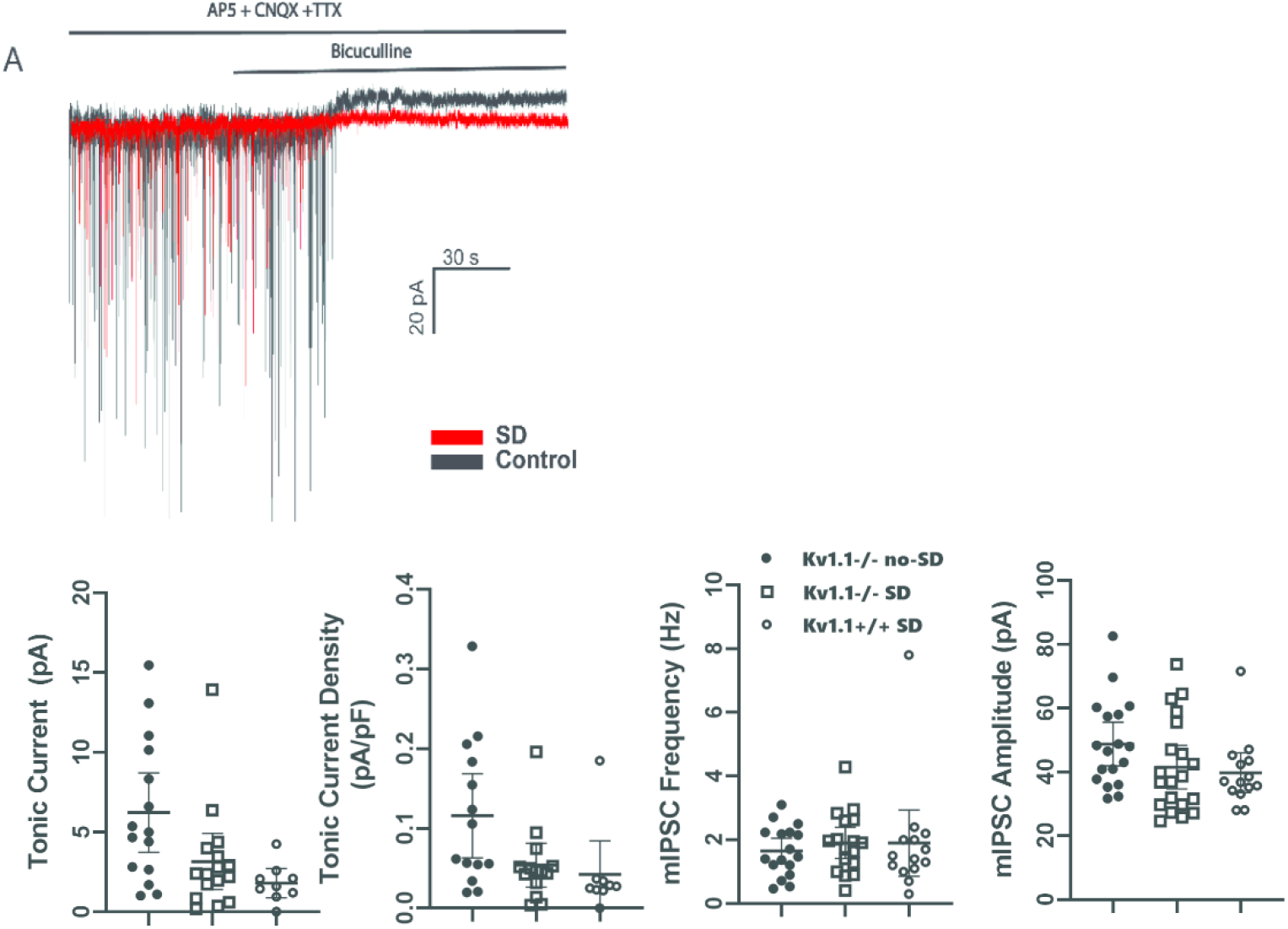
Data of a tonic current experiment is shown for SD and non-SD animals. **Panel A** shows tracing from a voltage clamp experiment in an SD and non-SD animal. Tracing shows mIPSCs after application of TTX (200nM), CNQX (10μM) and AP-5 (25μM) to block sodium channels, NMDA and AMPA/kainite receptors, respectively. Then, following addition of Bicuculline, mIPSCs were abolished and a shift on the holding current is seen which is attenuated in a SD animal. **Panel B & C:** The mean tonic inhibitory currents were 6.23±1.15 pA (±SEM) (15 cells from 4 animals) in the no-SD Kv1.1^-/-^ mice, 3.16±0.82 pA (16 cells from 6 animals) in Kv1.1^-/-^ with SD and 1.80±1.19pA in the wild-types with SD (9 cells from 5 animals) (ANOVA: F(2, 37)=5.238; *p*=0.009). The mean tonic current density (pA/pF) was 0.116±0.09 in no-SD Kv1.1^-/-^ whereas it was 0.054±0.04 in SD Kv1.1^-/-^ and 0.042±0.05 in SD wild types (ANOVA: F(2 35)=4.118, *p*=0.02) (means and 95% CI were shown). Post-hoc Tukey test showed that no-SD animals had much higher tonic current or current density that any SD group (p<0.05). **Panels D & E:** Mean mIPSC frequency (1.64±0.17 vs 1.89±0.19 vs 1.89±0.20 Hz) or amplitude (48.81±3.23 vs. 41.56±3.26 vs 39.75±2.89 pA) between any SD and non-SD groups is shown (means and 95% CI are shown) (*p*>0.05).

## Discussion

Our study demonstrates that SD in the Kv1.1^-/-^ model indeed exacerbates seizures. There was a nearly 52% mortality in the SD group that occurred between P37 and 44 whereas none occurred in non-SD controls, suggesting that SD superimposed on a condition where is sleep is already disrupted may affect survival. Interestingly, the effects of SD on seizures were lost after the 3^rd^ day of SD likely due to yet unknown homeostatic compensatory mechanisms, reminiscent of our prior observation where the adverse consequences of sleep fragmentation on memory function faded after the third day^10^.

Mechanistically, in a normal brain, SD is known to *decrease* excitatory synaptic transmission in the temporal lobe with altered expression of NMDA, AMPA receptors or their ratio^11^. In Transcranial Magnetic Stimulation (TMS) studies, SD reduced intracortical inhibition and increased excitability in focal or generalized epilepsy^12^. SD also increased susceptibility to seizures in penicillin, amygdala kindling and electroconvulsive shock models in cats, but the mechanisms are unknown^13^. However, there is a dearth of studies examining changes in excitatory or inhibitory synaptic transmission and cellular or network excitability with SD in epilepsy.

GABAergic signaling is the principal mechanism of inhibition in the CNS and consists of phasic and tonic inhibition, the former mediated by the release of presynaptic GABA that activates GABA_A_ and GABA_B_ receptors within the postsynaptic and peri-synaptic membrane whereas the latter is mediated by low ambient GABA that diffuses throughout the extracellular space^14^. GTI is mediated by GABA_A_ receptors containing α-4 and δ-subunits in the dentate gyrus and by receptors containing α-5 and γ subunits in the CA1 region of hippocampus^15^. Under baseline conditions *in vitro*, the total charge transferred by tonic current is considerably larger than that by phasic current, and thus, reduction in GTI is expected to increase overall excitability^16^. In models of temporal lobe epilepsy (TLE) there is evidence of reduction in GABA_A_ receptor α-5 (in CA1) and δ-subunits in dentate gyrus^17^, though surprisingly, several studies showed that tonic currents are normal or even enhanced in the hippocampus in mouse models^18^ or in human hippocampal tissue^19^. Interestingly however, SD diminished α-5 and δ-subunit containing GABA_A_ receptors in the hippocampus and diminished GTI in normal C57/BL6 mice^20^. Ganaxolone, a positive allosteric modulator of δ-subunit containing GABA_A_ receptors improved sleep in Kv1.1^-/-^ mice and enhanced survival though not significantly^20^. Diminished GTI may have contributed to increased excitability, resulting in SD-induced seizure exacerbation in Kv1.1^-/-^ mice. Thus, drugs that enhance GTI may have a therapeutic potential against SD-induced seizure exacerbation and needs further exploration. Overall, our results suggest that SD contributes to seizures acutely and leads to reductions in GTI that may also be relevant for epilepsy related mortality or SUDEP in humans.

Future studies are warranted to understand how the overall excitation-inhibition balance changes with SD. Mechanisms of diminished GTI from SD may include alterations in GABA_A_ receptor expression, variations in extracellular GABA concentrations or changes in endogenous neurosteroids that allosterically modulate receptors involved in GTI^21^. Regardless of the mechanism, sleep is already compromised in epilepsy due to seizures and/or associated sleep disorders, and any superimposed SD may have adverse consequences as demonstrated by the acutely exacerbated seizure burden and hastened mortality in our model.

## Acknowledgements

This work was funded by National Institutes of Health grant number R21NS104612-01A1 (PI:RM).

## Author contributions

1) Study concept and design: RM and MVJ. 2) Data acquisition: SSK, YP, EW, JAP did all experiments. 3) Analysis of data: RM, YP. 4) Drafting a significant portion of the manuscript or figures: RM, YP, EW, JAP, MVJ).

## Abbreviations

SD: Sleep deprivation
GTI: GABAergic tonic inhibition
GABA: gamma amino butyric acid
SUDEP: Sudden unexplained death in epilepsy
EEG: Electroencephalography
EMG: Electromyography
aCSF: artificial cerebrospinal fluid
NMDA: N-Methyl-D-aspartate
AMPA: α-amino-3-hydroxy-5-methyl-4-isoxazolepropionic acid
TLE: Temporal lobe epilepsy

